# Antifeedant activities of L-arabinose to caterpillars of the cotton bollworm *Helicoverpa armigera* (Hübner)

**DOI:** 10.1101/2020.07.20.213033

**Authors:** Longlong Sun, Zhenzhen Hong, Ying Ma, Wenhua Hou, Long Liu, Xincheng Zhao, Fengming Yan, Xinming Yin, Chenzhu Wang, Qingbo Tang

## Abstract

Exploring botanical biopesticide is one of the eco-friendly approaches for pest control in current crop protection. L-arabinose, a plant-specific and less absorptive pentose, is known for its selective inhibitory effect on the sucrose digestion in mammals. In this study, we investigated the effects of L-arabinose on the feeding preference, the peripheral gustatory perception, the larval development, as well as the activity of intestinal sucrase of an insect pest species, the cotton bollworm *Helicoverpa armigera* (Hübner) (Lepidoptera: Noctuidae), in an attempt to explore the antifeedant activities of this pentose. The results showed that L-arabinose deterred the feeding preferences of *H. armigera* caterpillars for its two host plants and two phagostimulants, the tobacco, the pepper, the sucrose and the fructose. Gustatory receptor neurons (GRNs) sensitive to L-arabinose were not found in the peripheral maxillary sensilla of *H. armigera* caterpillars, but the sensitivities of GRNs sensitive to sucrose, fructose and tobacco saps were suppressed by the additions of L-arabinose. Exposure of *H. armigera* caterpillars to dietary L-arabinose resulted in a prolonged larval developmental duration, a suppressed activity of intestinal sucrase and a reduced glucose level in midgut. *In vitro*, L-arabinose inhibited activities of the intestinal sucrase of *H. armigera* caterpillars in an uncompetitive manner. Taken together, these findings demonstrate that L-arabinose is a behavioral, gustatory and physiological inhibitor to caterpillars of the extremely generalist pest species *H. armigera*, suggesting the great potential of L-arabinose to be an effective antifeedant.

## Introduction

It is estimated that the insect pests are responsible for damaging between 20 and 40% of global crop production annually (FAO 2019). Though the strategy of the integrated pest management (IPM) has been extensively accepted in the past decades, the use of conventional synthetic insecticides is still the main approach for pest control in current crop protection. In comparison with conventional insecticides, many attentions have been paid to the behavioral manipulation of insect pests for their management in recent years (Dancewicz et al. 2020; Karkanis and Athanassiou 2021; Koul 2005; Walia et al. 2017; Wroblewska-Kurdyk et al. 2020). Of particular interests are nontoxic or less-toxic substances of natural origin that show selective activities toward insect pests but not toward non-target beneficial organisms and the environment (Akhtar et al. 2010; Junhirun et al. 2018; Koul et al. 2003; Ponsankar et al. 2020).

Most natural substances with behavioral manipulating activities could be perceived by the chemosensory system of insect pests, resulting in appetitive feeding behaviors or aversive feeding behaviors(Jiang et al. 2020; Mitri et al. 2009; Sollai et al. 2017; Sollai et al. 2015; Wada-Katsumata et al. 2013). Examples of manipulating activities of insect pests include repellent, antifeedant, deterrent, as well as growth inhibition and negative effects on both fecundity and longevity of the target species (Dancewicz et al. 2020; Hasheminia et al. 2013; Karkanis and Athanassiou 2021; Koul et al. 2003; Mao et al. 2019). Lots of substances originated from plants such as azadirachtin, rotenone, matrine, nicotine, pyrethrins have been well developed and commercially registered in some countries (Cheng et al. 2020; Isman 2020; Pavela 2016; Siegwart et al. 2015; Walia et al. 2017). But there should still be more botanical substances deserved to be explored for biological control of insect pests.

L-arabinose (abbreviated as LA), a naturally-occurring and widely present pentose, is a plant-specific sugar accounting for 5-10% of cell wall saccharides in varieties of plants including corn, rice, wheat, barley, rye, and oats (Bengtsson 1990; Garleb et al. 1989; Saulnier et al. 1995). LA is also known as a less absorptive sugar with a sweet taste and has been recognized to have a selective inhibitory effect on the intestinal sucrase activity in mammals, thereby resulting in a delay of sucrose digestion (Halschoujensen et al. 2015; Seri et al. 1996; Shibanuma and Houda 2011; Yamamoto 2013). In recent years, LA is widely used as additives in human food products with the intention to combat obesity and diabetes (DuBois and Prakash 2012; van Opstal et al. 2019).

In regard of inhibiting activity, we wonder whether LA has inhibitory effects on the digestion of sucrose in insect pests, and thereby LA may have the potential to be a phytochemical antifeedant. A pioneering report in 2010 found that the exposure of the sweet potato whitefly *Bemisia tabaci* to dietary arabinose resulted in low survivals of the nymphs and reduced mean day survivals of adults, suggesting a possible antifeedant activity of arabinose on this piercing insect species (Hu et al. 2010), but the proofs of how LA modulate the chemosensory perception and the feeding behaviors of insects were absent.

The cotton bollworm *Helicoverpa armigera* (Hübner) (Lepidoptera: Noctuidae) is an extreme generalist agricultural pest in the world, feeding on at least 161 host plant species in 49 plant families (Fitt 1989; Zalucki et al. 1986). In our preliminary studies, we observed that the feedings of *H. armgiera* caterpillars could be deterred by LA. Then we postulated that LA might be an antifeedant to this herbivorous chewing mouthpart pest species. In this study, we investigated the effects of LA on the feeding behavioral, the peripheral gustatory perception, the larval development, as well as the activities of intestinal sucrase both in *vitro* and in *vivo*, with the aim of exploring the antifeedant effects and the possible mechanism of this pentose to the lepidopteran pest species.

## Materials and Methods

### Insects

The colony of *H. armigera* was maintained in the laboratory under conditions of photoperiod (L16:D8), temperature (27 ± 1°C) and 75% relative humidity. Larvae of were reared on an standard artificial diet (Jiang et al. 2010) or standard artificial diet containing different concentrations of LA (Sigma-Aldrich) described as below. Adults were supplied with a 10% v/v solution of sucrose in distilled water with vitamins.

### Feeding choice assay

Six behavioral experiments were set to investigate the effects of LA on the feeding preferences of *H. armigera* caterpillars for plant leaves treated by different concentrations of LA (Table 1). Leaves from two plant species, the pepper *Capsicum frutescens* L. “Yu-Yi” and the tobacco *Nicotiana tabacum* “K326”, were used to be the feeding mediums respectively, in an attempt to investigate whether the preferences may change if different medium leaves were used. Of the 6 experimental settings, experiment II, IV and VI were used to investigate whether LA could counteract the stimulation effects of sucrose and fructose after LA was mixed to sucrose or fructose. While experiment III was used to investigate whether caterpillars more prefer sucrose if the control leaves were treated by sucrose.

**Table 1.** Settings of the dual-choice feeding assay of *H. armigera* caterpillars for LA-treated plant leaves. Note: The tobacco leaves and the pepper leaves were used as the medium leaf discs, respectively. LA: L-arabinose, Su.: sucrose; Fru.: fructose. Note in experiment II, III, IV and VI, leaves were treated by mixtures of LA and other sugar.

| Medium discs |  | Tobacco leaf |  |  |  | Pepper leaf |  |
| --- | --- | --- | --- | --- | --- | --- | --- |
| Experiments |  | I | II | III | IV | V | VI |
| Treatments | | LA ($10^{-4}$, $10^{-3}$, $10^{-2}$, $10^{-1}$, 1.0, 10, 67, 133, 200, 333 mM) | 200 mM Su. + (0, 100, 300, 400, 500, 600, 800 mM) LA | 10 mM Su. + (1.0, 10, 100, 200, 300, 400 mM) LA | 200 mM Fru. + (0, 300, 400, 500, 600 mM) LA | LA ($10^{-2}$, $10^{-1}$, 1.0, 10, 100, 200, 300, 400 mM) | 200 mM Su. + (0, 100, 300, 500, 600, 800, 1000 mM) LA |
| Control |  | pure water | pure water | 10 mM sucrose | pure water | pure water | pure water |

A dual-choice bioassay (Wang et al. 2017) was performed to test feeding preferences of the 5^th^ instar *H. armigera* caterpillars for leaf discs. In general, leaf discs (10 mm diameter) were punched from fresh plant leaves using a conventional hand-held punch, followed by immersed in a treatment solution or in a control solution (as shown in Table 1) for 5 min. For a test of a single caterpillar, a moist medium-speed qualitative filter paper (Ф11cm, Jiaojie, China) was put on the bottom of a Petri dish (12 cm diameter).Then four leaf discs treated with a treatment solution (A) and four discs with the control solution (B) were put on the moist filter paperin an ABABABAB fashion around the circumference. At test initiation, a single caterpillar was placed in the middle of the dish and the leaf disc consumption was observed at regular intervals until at least two of the four disks of either treatment (A) or control (B) had been consumed. Remaining areas of all discs consumed were measured using transparency film (PP2910, 3M Corp.). The feeding preference index was calculated as follows.

Preference index= (area of treatment-disc consumed-area of control-disc consumed) / (area consumed of all discs)

For each setting of choice scenario, the mean consumption of all caterpillars was calculated.

### Electrophysiological recordings

Four electrophysiological experiments were set to investigate the effects of LA on the sensitivity of gustatory receptor neurons (GRNs) in the styloconic sensilla located on the galea of 5^th^ instar *H. armigera* larvae by using the single sensillum recording (Roessingh et al. 1999; van Loon 1990). As shown in Table 2, experiment I and II investigated whether sensitive GNRs to LA were in the lateral sensillum and the medial sensillum. Experiment III investigated whether LA have effects on the sensitivities of the sucrose-best GRN in the lateral sensillum to sucrose. Experiment IV investigated whether LA have effects on the sensitivities of the GRNs sensitive to tobacco leaf saps in the medial sensillum. The sucrose best GRN in the lateral sensillum and the GRNs sensitive to tobacco-leaf-sap best in the medial sensillum have been identified in our previous work (Ma et al. 2016; Tang et al. 2014). The leaf saps were extracted as per our previous work (Tang et al. 2014).

**Table 2.**
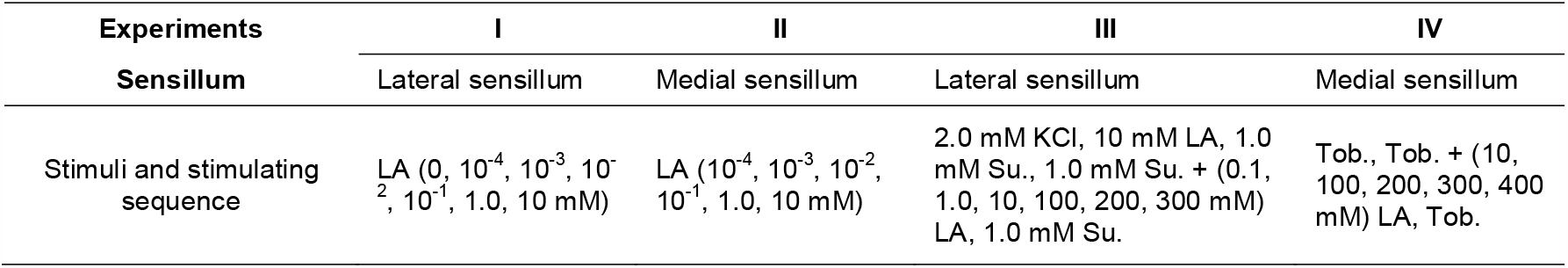
The substances with different concentrations used in the electrophysiological recordings. Note: LA: L-arabinose, Su.: sucrose; Tob.: tobacco leaf saps. 2 mM KCl was the adequate electrolyte solvent.

Before stimulation, an excised 5^th^ instar caterpillar head was mounted on a silver wire electrode, which was connected to the input of a pre-amplifier (Syntech Taste Probe DTP-1, Hilversum, The Netherlands). Glass micropipette was filled with stimulating solution and touched the tip of a sensillum under the control of a micromanipulator. The stimuli and the stimulating sequence in a single caterpillar were shown in Table 2. Amplified signals were digitized by an A/D-interface (Syntech IDAC-4) and sampled into a personal computer.

The analysis of responding spikes to different stimuli were performed with the aid of Autospike version 3.7 software (Syntech, The Netherlands). In general, impulse waveforms were classified into different groups according to the amplitude, waveform (shape), and phasic temporal pattern, as well as by visual inspection, and then were counted. In most recordings, three or four impulse spikes were identified and labeled as small (S), intermediate (M1 or/and M2), and large (L), which best responded to water, phagostimulant/deterrent, and salt, respectively (Sollai et al. 2014; Sun et al. 2021).The mean impulse frequency (spk.s^-1^) of a GRN to one stimulus in the first 1000 ms was calculated.

### Effects of LA on Larval development

Six groups of *H. armigera* neonatal caterpillars were reared by a standard artificial diet (Jiang et al. 2010) containing different concentration of LA from 0, 6.66 mM, 66.6 mM, 199.8 mM, 333 mM, to 500 mM until the last instar, respectively. The larval developmental duration and the mortality of caterpillars were recorded. Caterpillars were maintained through pupation to adult emergence, and then the pupal developmental duration as well as the pupal weights was recorded.

### Effects of LA on enzymatic activity

Six groups of neonate caterpillars of *H. armigera* were reared by standard artificial diets containing different concentrations of LA from 0, 6.66 mM, 66.6 mM, 199.8 mM, 333 mM, to 500 mM until the second day of the 5^th^ instar. Then the midguts of caterpillars were extracted and weighted. Midguts were hand homogenized in PBS (pH 7.4) at a ratio of 1.0 g midgut to 6.0 ml PBS, and then were centrifuged at 2500 g for 10 min at 4°C. Supernatants were used to analyze sucrase activity, sucrose concentration, and glucose concentration by manufacturer protocols (No. A082-2 for sucrase activity, No.A099 for sucrose, and No.F006 for glucose; Nanjing Jiancheng Bioengineering Institute, Nanjing, China).

### Kinetic analysis of sucrase inhibition by LA

Midguts of the fifth instar caterpillars of *H. armigera* were dissected and then were rinsed with ice-cold saline (10%) followed by weighing and hand-homogenization. Homogenized tissues were centrifuged at 2500 rpm for 10 min at 4°C. Supernatants were transferred to a fresh tube and stored at −20°C until use.

For the kinetic studies of the inhibitory effects of LA on sucrase activity in caterpillars, midgut supernatants prepared as above were incubated with increasing concentrations of sucrose (0, 10 mM, 20 mM, 30 mM, 40 mM, 50 mM, 60 mM) in the absence or presence of LA (6.66 mM or 9.99 mM) or acarbose (7.35 μM or 11.79 μM; Sigma-Aldrich). Selected concentrations of these sugars were determined based on preliminary experiments.

The volume of standard reaction assay was 90 μL. After the reaction mixtures incubated at 37°C for 5 min, reactions were terminated by incubating in boiling water for 5 min. After cooldown, the D-glucose concentration was measured using a Glucose Oxidase Method (No. F006; Nanjing Jiangcheng Bioengineering Company, Nanjing, China). Specific activity was calculated as micromoles of substrate hydrolyzed per milligram protein per min. Each assay was repeated at least three times and results were analyzed by Lineweaver-Burk plots.

### Data analysis and statistics

All data are presented as mean ± SEM. A one-sample *t*-test was used to compare the means of the preference indices of caterpillars between for control leaves and for treatment leaves (*P* < 0.05). The levels of significance were as follows: NS (non-significant): *P* ≥ 0.05, *: *P* < 0.05, **: *P* < 0.01, ***: *P* < 0.001. For statistical analysis of neuronal responding frequency to stimuli, the original responding frequencies were square-root transformed if the variances were not equal. A one-way ANOVA followed by a Scheffé test was used to compare the mean responding frequency of the same GRN to stimuli with different concentrations (*P* < 0.05).

The data of the developmental parameters and the midgut enzyme activities were log-transformed for uneven sample sizes. Means were compared by one-way ANOVA followed by a Scheffé Test (*P* < 0.05). All statistical analyses were performed using SPSS version 10.0 (SPSS Inc., Chicago, USA).

## Results

### Inhibition of LA on feeding behaviors

The results of the feeding preference of the 5^th^ instar caterpillars of *H. armigera* for LA treated tobacco leaves and for pure distilled water treated tobacco leaves were shown in figure 1. It showed that low concentrations of LA from 0.0001 to1.0 mM had no significant effects on the preferences of caterpillars (Fig. 1A: 0.0001mM, *t*(107)= 0.409, *P =* 0.686; 0.001mM, *t*(86) = −0.935, *P =* 0.352; 0.01mM, *t*(160)= −1.157, *P =* 0.249; 0. 1mM, *t*(117) = −1.319, *P =* 0.190; 1.0 mM, *t*(224) = −1.679, *P =* 0.095). But high concentrations of LA ≥ 10 mM significantly deterred the feeding of caterpillars (Fig. 1A: 10mM, *t*(272) = −2.77, *P =* 0.006; 66.6mM, *t*(243) = −3.491, *P =* 0.001; 133mM, *t*(190) = −5.88, *P* < 0.0001; 200mM, *t*(311) = −6.359, *P* < 0.0001; and 333mM, *t*(100) = −5.743, *P* < 0.0001).

**Figure 1.**
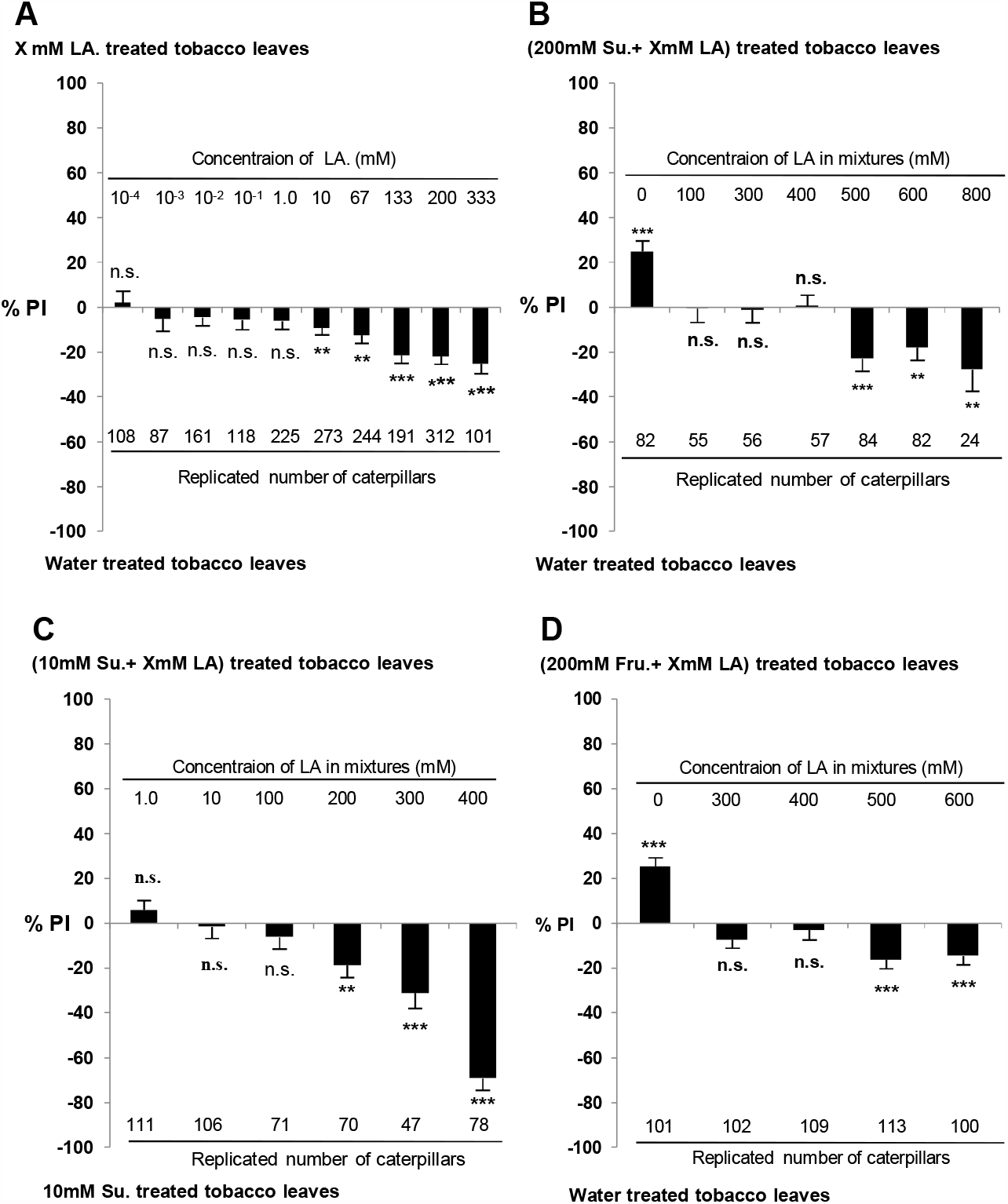
Effects of LA on the feeding preferences of *H. armigera* caterpillars for tobacco leaves. Columns represent the mean preference indices ± SEM of the 5^th^ instar caterpillars for tobacco leaves treated with LA/mixtures or water (control) in dual-choice assays. **A:** Preferences between leaves treated by LA with different concentrations and water-treated leaves (control). **B:** Preferences between water-treated leaves (control) and leaves treated by mixtures of 200 mM sucrose (Su.) plus LA with different concentrations. **C:** Preferences between 10 mM sucrose-treated leaves and leaves treated by mixtures of 10 mM sucrose plus LA with different concentrations. **D:** Preferences between water-treated leaves (control) and leaves treated by mixtures of 200 mM fructose (Fru.) plus LA with different concentrations. A one-sample t-test was used to compare the means of the preference indices for treatment leaves and for control leaves in each experimental setting (*P* < 0.05).

We also found LA could counteract the phagostimulant effects of sucrose on *H. armigera* caterpillars. Caterpillars significantly preferred 200 mM sucrose-treated leaves over control leaves (*t* (81) = 5.117, *P* < 0.0001), whereas caterpillars preferred similarly to control leaves and leaves treated by mixtures of 200 mM sucrose with 100, 300, and 400 mM LA, respectively (200mM sucrose + 100mM LA, *t*(154) = −0.048, *P =* 0.962; 200mM sucrose +300mM LA, *t*(55) = −0.156, *P =* 0.877; 200mM sucrose +400mM LA, *t*(56) = 0.138, *P =* 0.891) (Fig. 1B). With the concentration of LA increasing in mixtures, caterpillars exhibited aversive feedings to mixture-treated leaves (200mM sucrose + 500 mM LA, *t*(83) = −4.224, *P =* 0.0001; 200 mM sucrose + 600 mM LA, *t*(81) = −3.118, *P =* 0.0025; 200 mM sucrose + 800 mM LA, *t*(23) = −2.876, *P =* 0.0085) (Fig. 1B).

Next, we examined whether *H. armigera* caterpillars could distinguish between tobacco leaves treated by 10 mM sucrose and leaves treated by mixtures of 10 mM sucrose with different concentrations of LA. It showed that low concentrations of LA from 1.0 mM to 100 mM in mixtures had no significant effects on the preferences (10 mM sucrose + 1mM LA, *t*(110) = 1.363, *P =* 0.176; 10 mM sucrose + 10mM LA, *t* (105) = −0.301, *P =* 0.764; 10 mM sucrose + 100mM LA, *t*(70) = −1.121, *P =* 0.266), but leaves treated by mixtures containing LA ≥ 200 mM drove significant aversive feedings of caterpillars (10 mM sucrose + 200 mM LA, *t*(69) = −3.362, *P =* 0.001; 10 mM sucrose + 300 mM LA, *t*(46) = −4.396, *P* < 0.0001; 10 mM sucrose + 400 mM LA, *t*(77) = −13.09, *P* < 0.0001) (Fig. 1C).

We also found that LA could counteract the stimulating effect of another phagostimulant, the fructose. Single 200 mM fructose significantly elicited appetitive feedings of *H. armigera* caterpillars (*t*(101) = 6.722, *P* < 0.0001), but mixtures containing 200 mM fructose with 300mM or 400mM LA induced a similar preferences as controls (200 mM fructose + 300 mM LA, *t*(102) = −1.911, *P =* 0.059; 200 mM fructose + 400 mM LA, *t*(109) = −0.677, *P =* 0.500) (Fig. 1D). When the concentration of LA in mixtures increased to 500 mM and 600 mM, mixture-treated leaves significantly drove aversive feedings of *H. armgiera* caterpillars (200 mM fructose+ 500 mM LA, *t*(113) = −4.115, *P =* 0.0001; 200 mM fructose + 600 mM LA, *t*(100) = −0.3.572, *P =* 0.0005) (Fig. 1D).

When the pepper leaves were used as the plant leaf medium, LA still showed inhibiting effects on feedings of *H. armigera* caterpillars. Caterpillars showed more and more aversive feedings for LA-treated pepper leaves with the concentration of LA increasing (Fig. 2A: 0.01mM LA, *t*(62) = 0.124, *P =* 0.902; 0.1mM LA, *t*(252) = −1.315, *P =* 0.190; 1.0 mM LA, *t*(214) = −1.493, *P =* 0.137; 10mM LA, *t*(325) = −2.251, *P =* 0.025; 100mM LA, *t*(155) = −2.028, *P =* 0.044; 200mM LA, *t*(115) = −2.244, *P =* 0.027; 300mM LA, *t*(95) = −5.211, *P* < 0.0001; 400mM LA, *t*(107) = −4.360, *P* < 0.0001).

**Figure 2.**
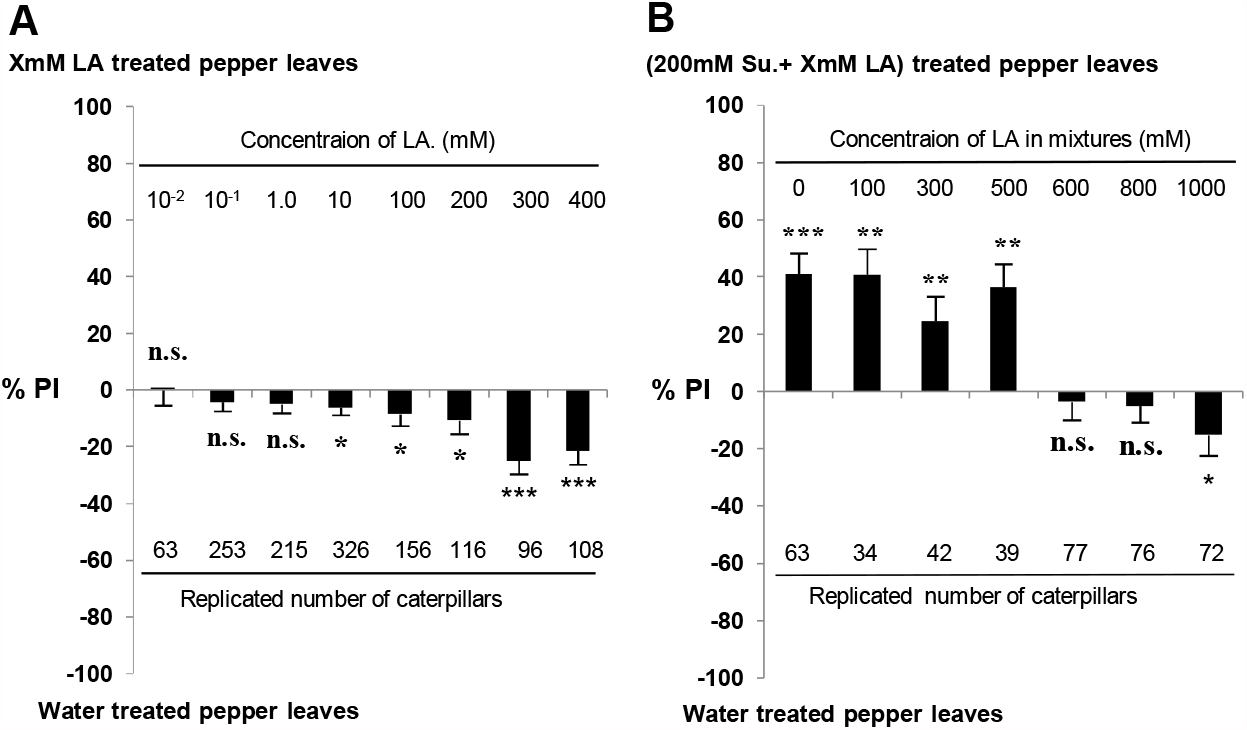
Effects of LA on the feeding preference of *H. armigera* caterpillars for pepper leaves. Columns represent the mean preference indices ± SEM of the 5^th^ instar caterpillars for pepper leaves treated with LA/mixtures or water (control) in dual-choice assays. **A:** Preference between for leaves treated by LA with different concentrations and for water-treated leaves (control). **B:** Preference between for water-treated pepper leaves (control) and for pepper leaves treated by mixtures of 200 mM sucrose (Su.) plus LA with different concentrations. A one-sample t-test was used to compare the means of the preference indices for treatment leaves and for control leave in each experimental setting (*P* < 0.05).

Similarly, LA also could counteract the phagostimulant effects of sucrose on *H. armigera* caterpillars when the pepper leaves were used as the medium leaves. Mixtures containing 200 mM sucrose with LA from 100 mM to 500 mM still induced the preference of caterpillars for sucrose (Fig. 2B: 200 mM sucrose + 0 LA, *t*(62) = 5.718, *P* < 0.0001; 200 mM sucrose + 100 mM LA, *t*(33) = 4.630, *P =* 0.0001; 200 mM sucrose + 300 mM LA, *t*(41) = 2.899, *P =* 0.0060; 200 mM sucrose + 500 mM LA, *t*(38) = 4.521, *P =* 0.0001). Whereas LA in mixtures ≥ 600 mM, the preferences for sucrose were significantly counteracted (Fig. 2B: 200 mM sucrose + 600 mM LA, *t*(76) =-0.564, *p* = 0.574; 200 mM sucrose + 800 mM LA, *t*(75) = −0.851, *p* = 0.398). When LA in mixtures increased to 1000 mM, caterpillars significantly preferred control leaves over mixture-treated leaves (Fig. 2B: 200 mM sucrose + 1000 mM LA, *t*(71) = −2.110, *p* = 0.038).

### Inhibition of LA on gustatory sensitivity

Gustatory perception in *H. armigera* caterpillars are mainly dominated by two pairs of gustatory styloconic sensilla, the lateral sensilla and the medial sensilla (Sun et al. 2021; Tang et al. 2014; Zhou et al. 2010), located on the maxillary galea of mouthparts (Figs. 3A and 3B). But our current results showed LA from 0.0001 mM to 10 mM stimulated very low responses, which is similar to those induced by 2 mM KCl in the response to the two styloconic sensilla (Fig. 3C), suggesting non-specific GRN housed in both the two sensilla were sensitive to LA.

**Figure 3.**
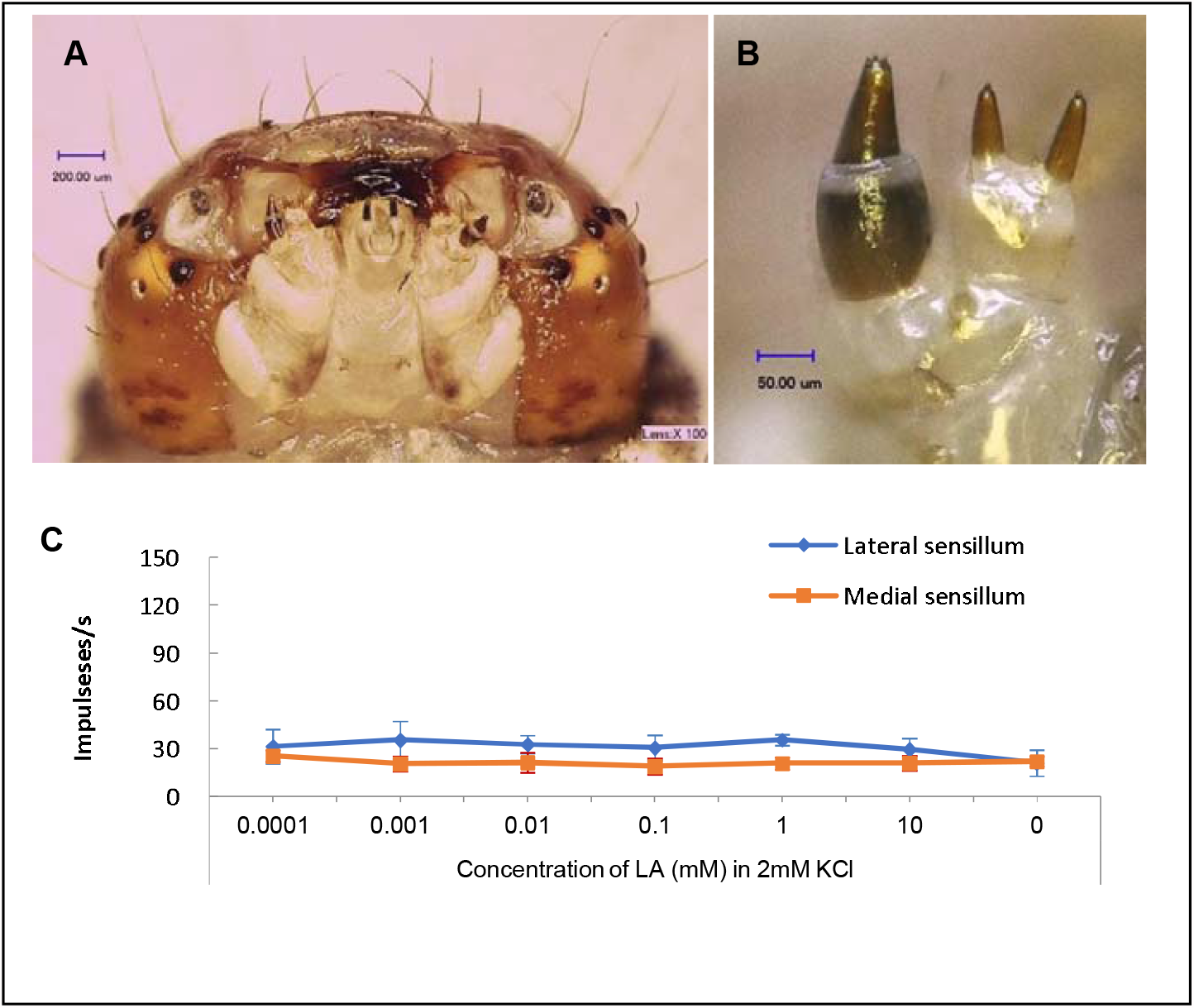
Morphology and electrophysiological response of gustatory sensilla of *H. armigera* caterpillars. **A:** Ventral view of the head showing the mouthparts of fifth instar caterpillars. **B:** Close-up view of the medial and the lateral sensillum styloconicum located on a galea. Images were taken by the digital microscope (VHX-600, Keyence, Japan). **C:** Responding curves of the lateral sensillum and the medial sensillum to series concentrations of LA in 2.0 mM KCl.

We then investigated whether LA affect the sensitivities of GRNs sensitive to other plant substances, such as the phagostimulant sucrose and the leaf saps from the host plant tobacco, by testing the responses of sensillum to mixtures containing LA and these substances. Firstly we tested the effects of LA on the sensitivity of the sucrose-best GRN in the lateral sensillum of the 5^th^ instar *H. armigera* caterpillars. The results showed that both 2.0 mM KCl and 10 mM LA only induced weak responses of a water-like L-GRN and a salt-like S-GRN, while 1.0 mM sucrose induced activities of three GRNs in the lateral sensillum of caterpillars, including the weak responses of L-GRN and S-GRN, and the strong response of the sucrose-best M1-GRN (Fig. 4A). But the sensitivity of the sucrose-best M1-GRN decreased with the concentration of LA increased in mixtures including 1.0 mM sucrose (ANOVA, *P* < 0.05). Even the sensitivities of the L-GRN and the M1-GRN were almost completely suppressed when the concentration of LA in mixtures reached to 200mM and 300mM (Fig. 4A, Fig. 4B). Finally, when the lateral sensillum was repeated stimulated by 1.0 mM sucrose in the absence of LA, the activity of the sucrose-best M1-GRN recovered completely (Figs. 4A, 4B).

**Figure 4.**
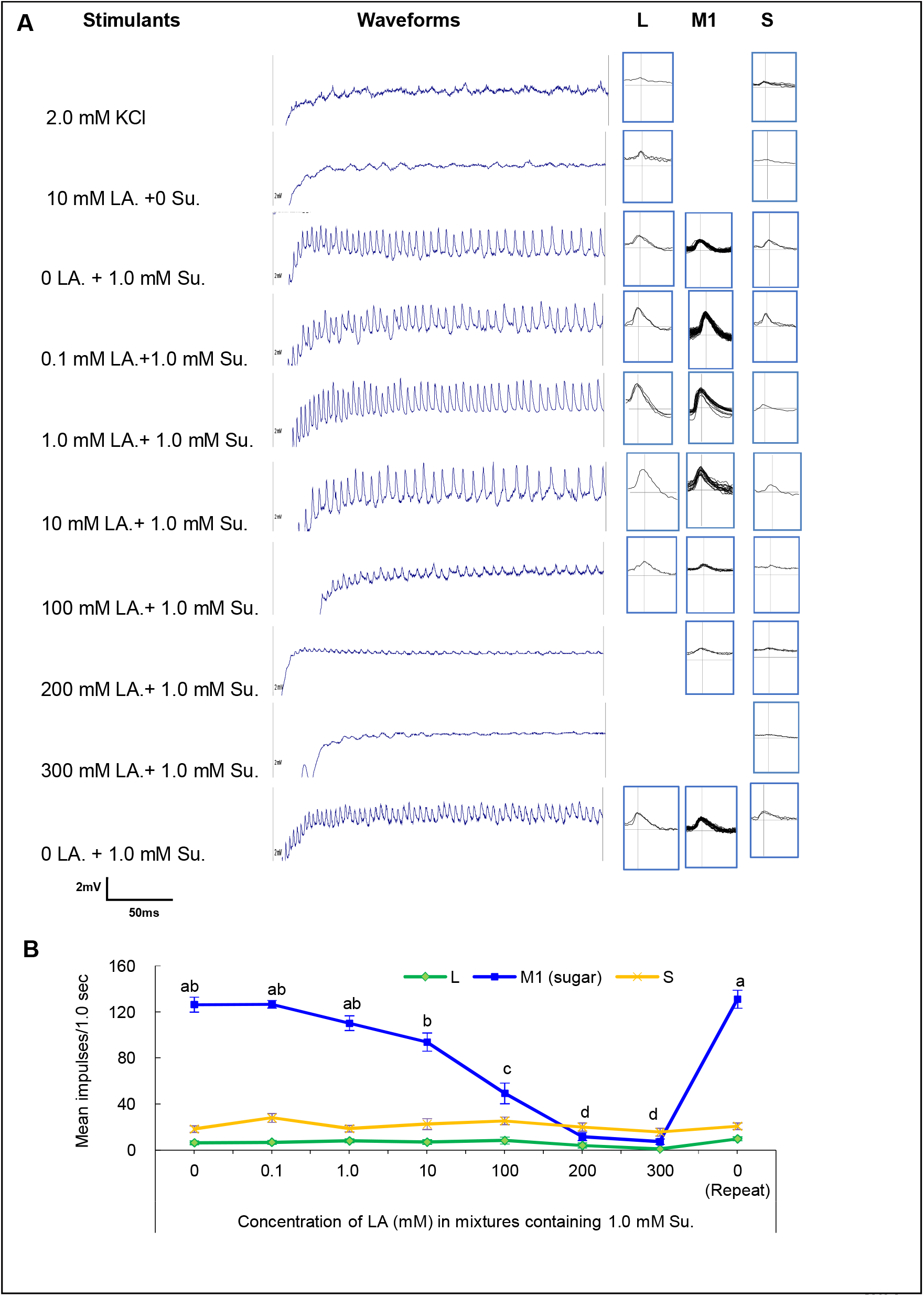
Representative responding traces and effects of LA on the sensitivity of the sucrose best GRN in the lateral sensillum. **A:** Representative traces (200 ms) and spikes identified from traces. Different traces were obtained from a single lateral sensillum in the same caterpillar to different stimuli with the stimulating order from up to down. L, M1, and S represent the identified spikes of GRNs sensitive to water, sucrose and 2mM KCl, respectively. **B:** Curves of responding frequency ± SEM of identified GRNs to 1.0 mM sucrose (Su.) and mixtures containing 1.0 mM sucrose and different concentrations of LA. The Scheffé’s post hoc test for one-way ANOVA was used to compare the mean response frequency of M1 to different stimuli (*P* < 0.05). Columns having no lowercase letters in common differ significantly (*P* < 0.05).

We also found that LA suppressed the sensitivities of GNRs in the medial sensillum of *H. armigera* caterpillars sensitive to tobacco leaf saps. As shown in figure 5, pure tobacco leaf saps induced the activities of 4 GRNs in the medial sensillum including the strong response of the M1-GRN, the relatively strong response of the M2-GNR, the weak responses of L-GNR and S-GNR, but the activities of the M1-GRN and the M2-GRN decreased with the concentration of LA in mixtures increasing. Even the activities of the M1-GRN, M2-GRN and L-GRN almost were completely inhibited when LA reached to 300 mM and 400 mM in mixtures (Figure 5A and 5B). Finally, when the medial sensillum was repeated stimulated by tobacco leaf saps in the absence of LA, the activity of the GRNs sensitive to tobacco leaves recovered completely (Figs. 5A, 5B).

**Figure 5.**
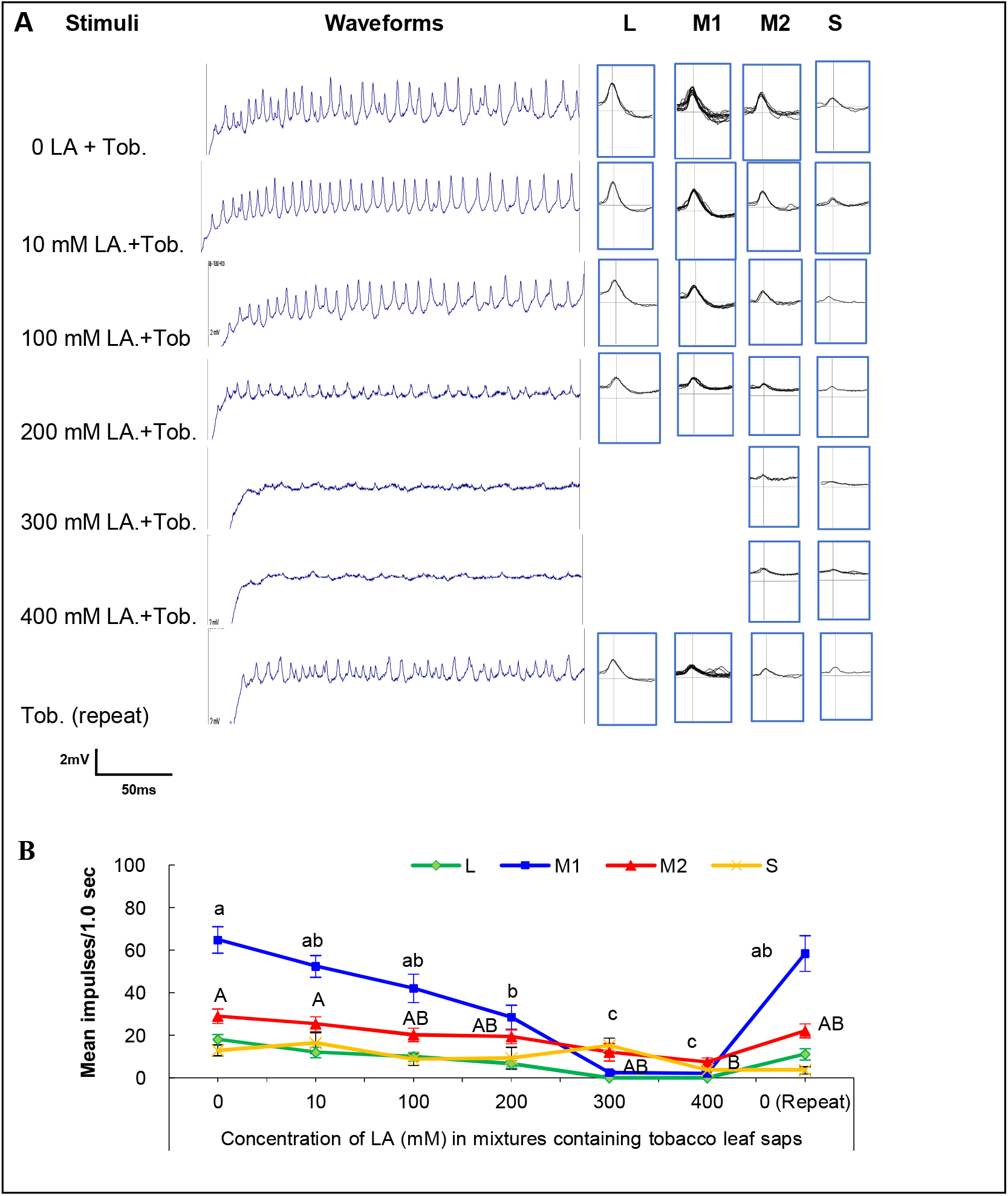
Representative responding traces and effects of LA on the sensitivities of GRNs sensitive to tobacco leaf saps in the medial sensillum. **A:** Representative traces (200 ms) and spikes identified from traces. Different traces were obtained from a single medial sensillum in the same caterpillar to different stimuli with the stimulating order from up to down. L, M1, M2, and S represent the identified spikes of GRNs sensitive to water, sugars, deterrents and 2mM KCl, respectively. **B:** Curves of responding frequency ± SEM of the identified GRNs to mixtures containing tobacco saps and different concentrations of LA. The Scheffé’s post hoc test for one-way ANOVA was used to compare the mean response frequency of M1 and M2 to different stimuli (*P* < 0.05). Columns having no capital letters or lowercase letters in common differ significantly *(P* < 0.05).

### Larval development prolonged after exposed to LA

The larval development duration, the larval mortality rate, the puapl development duration, and the pupal weight of *H. armgiera* originated form neonates reared on artificial diets containing serial concentrations of LA (6.66 mM, 66.6 mM, 199.8 mM, 333 mM, and 500 mM) were also examined. The results show that dietary LA has significant effects on the larval developmental duration, the pupal developmental duration and the pupal weights of *H. armigera* (one-way ANOVA: larval developmental duration, *F*(5, 473) = 48.928, *P* < 0.0001; pupal developmental duration, *F*(5, 379) = 5.384, *P =* 0.02; pupal weights, *F*(5, 466) = 11.125, *P* < 0.0001) (Fig. 6). At the concentrations of 333 mM and 500 mM of LA, the larval developmental duration was significantly prolonged compared to the control caterpillars (Scheffé Test: *P* < 0.001; Fig. 6A). The larval mortalities were not significantly affected by dietary LA (one-way ANOVA: *F*(5, 18) = 0.283, *P =* 0.917, Fig. 6B). But the pupal duration and the pupal weight were significantly decreased under 500 mM LA (Scheffé Test followed ANOVA: pupation duration duration, *P =* 0.005; pupal weights, *P* < 0.001; Fig. 6C and 6D). The developmental assays demonstrate that the consumption of LA by *H. armigera* caterpillars could result in the delaying of larval development.

**Figure 6.**
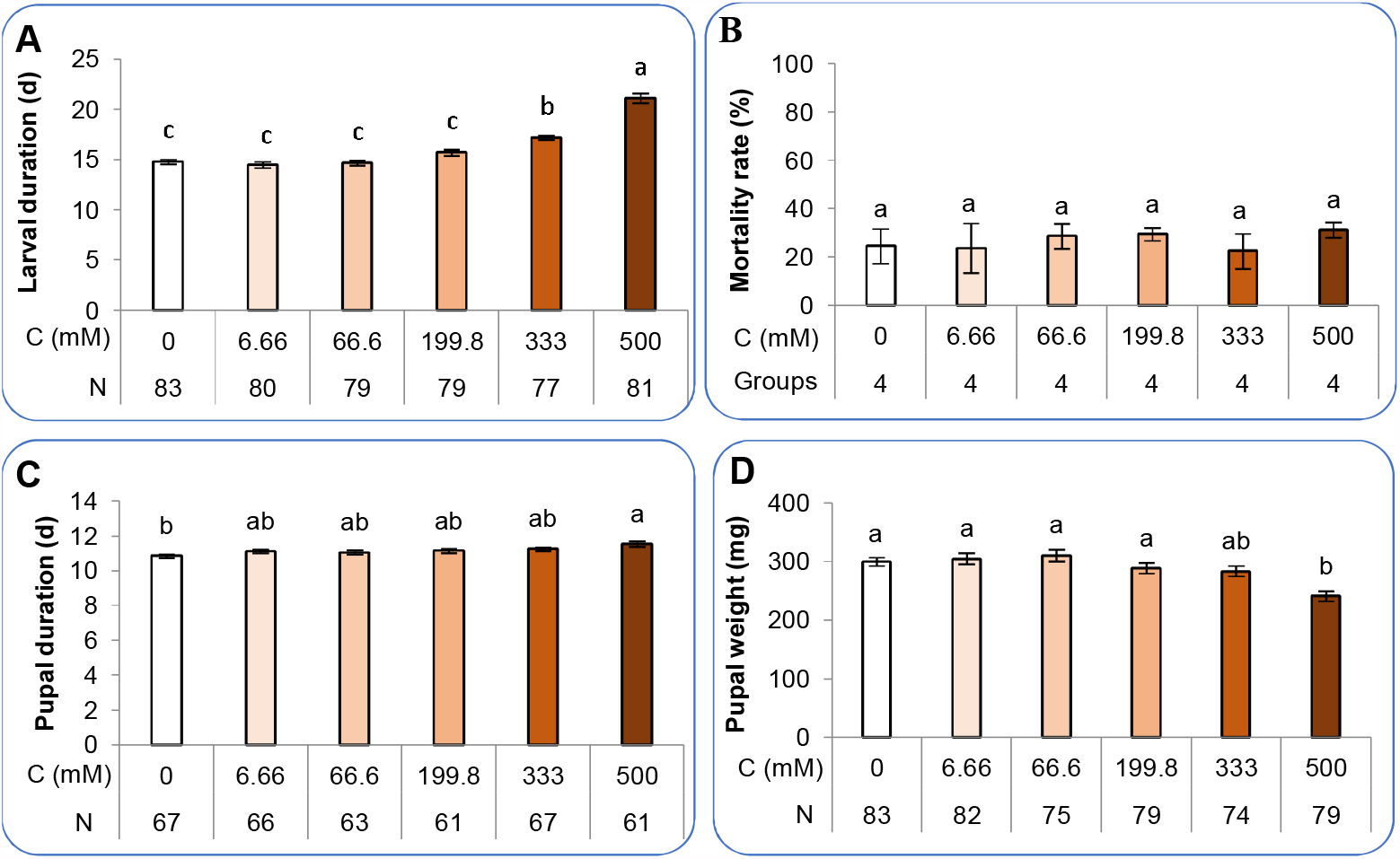
Effects of LA on the development of *H. armigera*. Comparison of the larval developmental duration (**A**) and the larval mortality rate (**B**) after feeding larvae on artificial diets containing different concentrations of LA from neonate until pupation. Comparison of pupal developmental duration (**C**) and pupal weight (**D**) of pupae originated from neonatal caterpillars rearing on artificial diets containing different concentrations of LA until pupation. The Scheffé’s post hoc test for one-way ANOVA was used to compare the means of developmental parameters of caterpillars reared on different dietary LA (*P* < 0.05). Columns having no lowercase letters in common differ significantly (*P* < 0.05). C: concentration of LA; N: the number of caterpillars tested. Groups: repeated numbers of the larval mortality were investigated.

### Suppressed activity of digestive enzymes after exposure to LA

After exposures of *H. armigera* neonates to dietary LA until the 5^th^ instar stage, the sucrase activity in midguts of the 5^th^ instar *H. armgiera* caterpillars were suppressed at a dose-dependent manner (6.66 mM – 500 mM; One-way ANOVA: *F*(5, 134) = 26.755, *P* < 0.0001) (Fig. 7A). The sucrase activities in midguts of caterpillars exposed to 6.66 mM and 66.6 mM dietary LA were similar as those of the control caterpillars (One-way ANOVA, *P =* 0.692), while the sucrase activities of caterpillars exposed to 199.8 mM, 333 mM and 500 mM dietary LA were significantly lower than those of caterpillars exposed to 6.66 mM and 66.6 mM dietary LA and the control caterpillars (one-way ANOVA: *P* < 0.01, all results; Fig. 7A).

**Figure 7.**
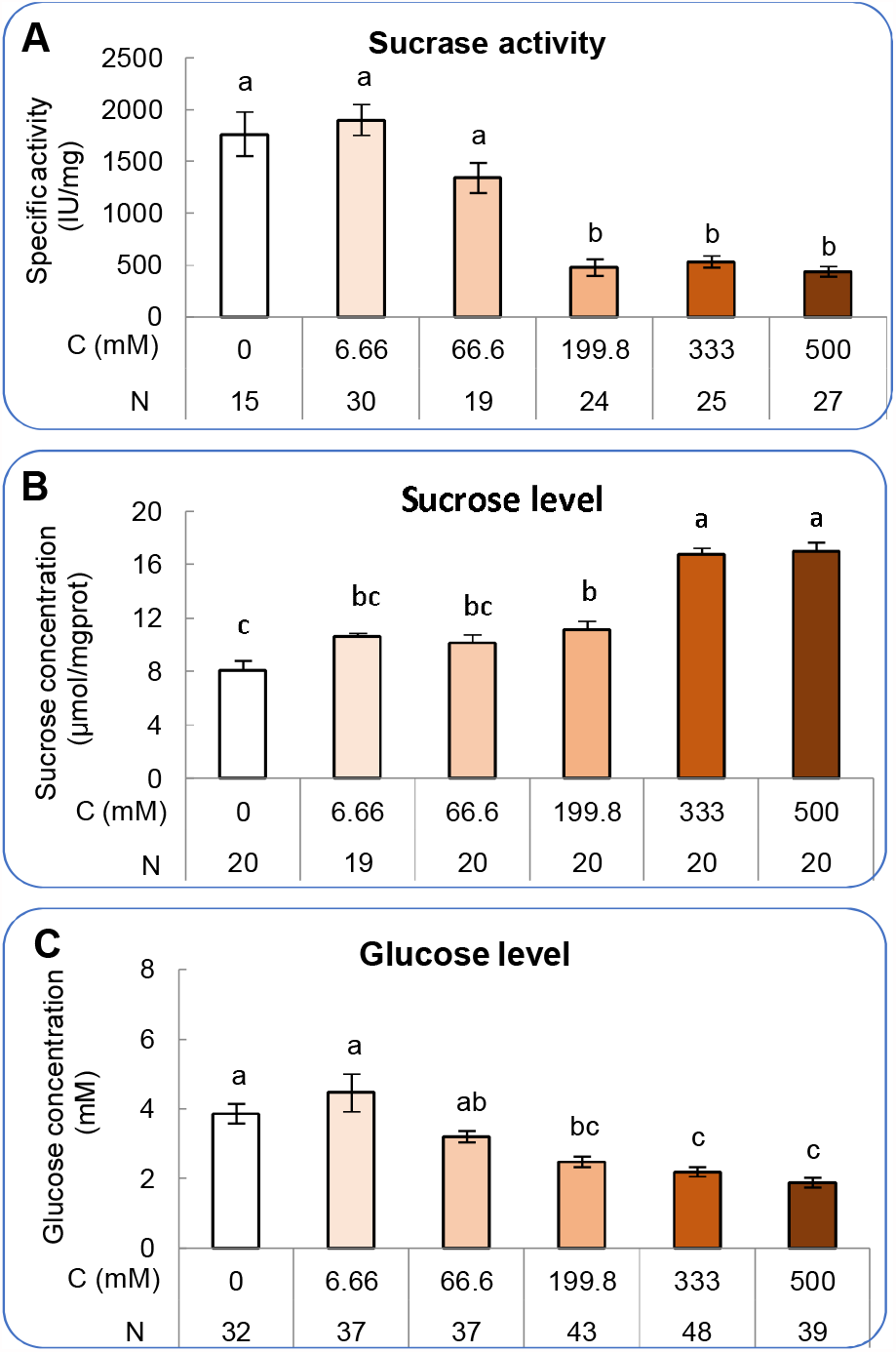
Effects of LA on the intestinal physiology of *H. armigera* caterpillars. Comparison of the sucrase activity (**A**), sucrose concentration (**B**), and the glucose concentrations (**C**) in midgut of the 5^th^ instar caterpillars exposed to artificial diets containing different concentrations of LA from neonate until pupation. The Scheffé’s post hoc test for one-way ANOVA was used to compare the means of physiological parameters of caterpillars exposing to different concentrations of LA (*P* < 0.05). Columns having no lowercase letters in common differ significantly (*P* < 0.05). C: concentration of LA; N: the number of caterpillars tested.

As expected, the sucrose concentration in midgut of caterpillars exposed to dietary LA increased at a dose-dependent manner (One-way ANOVA: *F*(5, 113) = 39.750, *P* < 0.0001) (Fig. 7B). The sucrose concentration of caterpillars exposed to 199.8 mM LA was significantly higher than that of the control caterpillars (Scheffé’s Tests: *P =* 0.023; Fig. 7B), but was similar to that of caterpillars exposed to 6.66 mM and 66.6 mM LA (Scheffé’s Tests: *P =* 0.926; Fig. 7B). With the dietary LA increasing to 333 mM and 500 mM, the sucrose concentration in midguts of caterpillars were significantly higher than that of caterpillars exposed to low concentrations of dietary LA and the control (Scheffé’s Tests: *P* < 0.0001, all results; Fig. 7B).

Correspondingly, the glucose levels in midgut of LA-fed caterpillars decreased in a dose-dependent manner from 6.66 mM to 500 mM dietary LA (One-way ANOVA, *F*(5, 230) = 17.590, *P* < 0.0001). Caterpillars exposed to 0, 6.66 mM and 66.6 mM dietary LA were similar at their midgut glucose levels (Scheffé’s Tests followed ANOVA: *P =* 0.08; Fig. 7C). But the glucose levels in caterpillars exposed to 199.8 mM dietary LA were significantly lower than that in control caterpillars and caterpillars exposed to 6.66 mM LA (Scheffé’s Tests: *P* < 0.01; Fig. 7C). With the concentration of LA in diets increasing to 333 mM and 500 mM, the glucose levels were even lower than that of caterpillars exposed to 66.6 mM dietary LA (Scheffé’s Tests: all *P* < 0.05; Fig. 7C).

### Kinetic analysis of sucraseinhibition by LA

We also found that LA inhibited the activity of intestinal sucrase of *H. armigera* caterpillars in an uncompetitive manner *in vitro* based on the Lineweaver-Burk analyses, displaying parallel lines of 1/*V* for the different LA concentrations (Fig. 8A). The maximum reaction velocity (*V*_max_) decreased from 7.31 to 5.85 mmol/min/mg protein and the *K*_m_ decreased from 7.24 to 5.15 mM with increasing of the LA concentration from 0 to 9.99 mM. At concentrations of 6.66 and 9.99 mM, LA reduced sucrase activity 12.24% and 26.81%, respectively, at *V*_max_ with the *K*_*i*_ of 30.66 mM/L.

**Figure 8.**
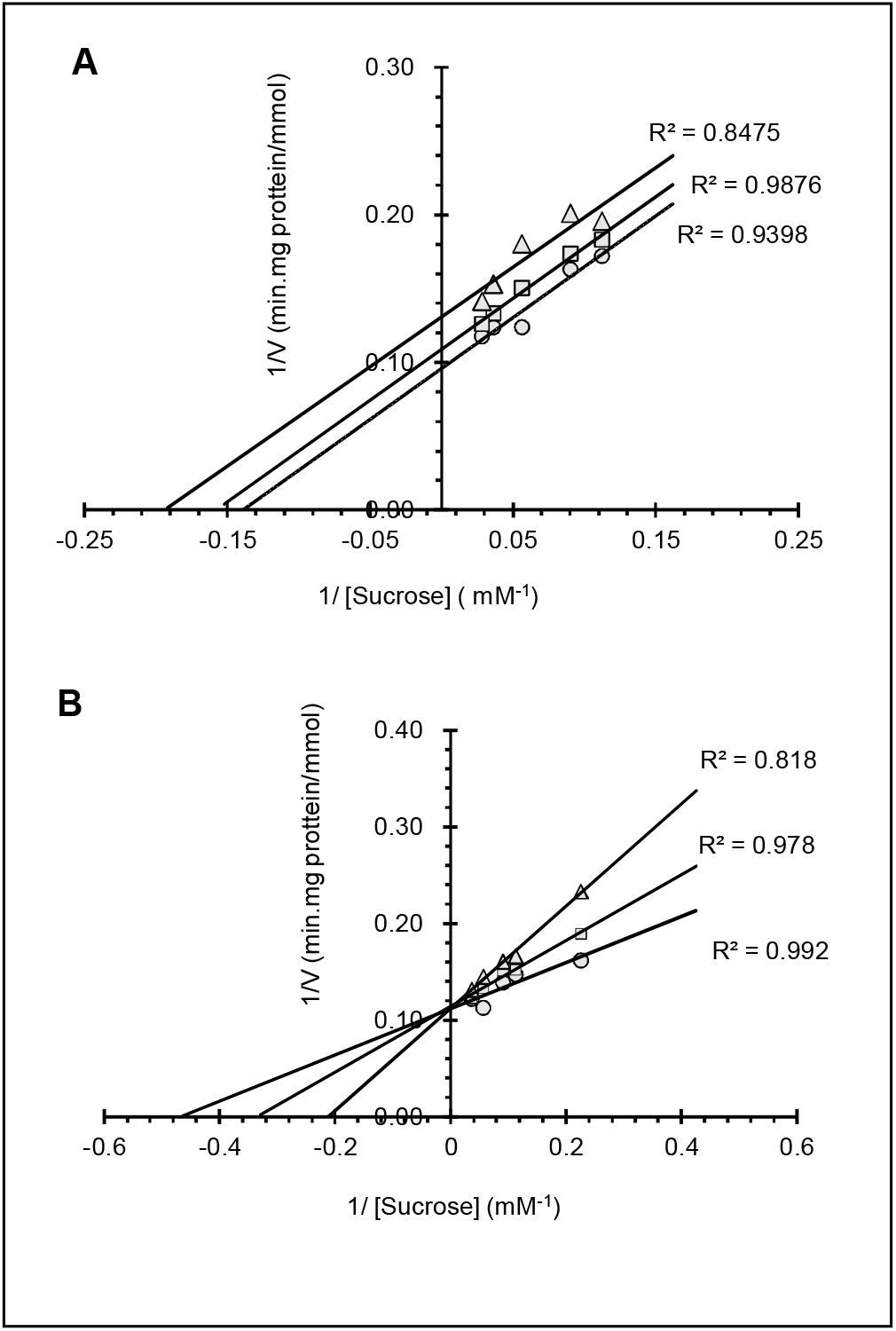
Kinetic analysis of sucrase inhibition by LA (A) and acarbose (B). Results represent the Lineweaver-Burk analysis. Supernatants extracted from midgut homogenates of the 5^th^ instar caterpillar were incubated with increasing concentrations of sucrose in the presence or absence of inhibitor: (**A**) 6.66 mM LA (□), 9.99 mM LA (□), and the control in the absence of LA (○); (**B**) 7.53 μM acarbose (□), 11.79 μM acarbose (□), and the control in the absence of acarbose (○). R^2^ was the coefficient of determination which predicts the goodness of fit for the linear regression model. *V* was the velocity of the reaction.

As the positive control, the other alpha-glucosidase inhibitor, acarbose, inhibited sucrase activities by a fully competitive manner (Fig. 8B). The *V*_max_ was similar from 8.94 to 8.89 mmol/min/mg protein and the *K*_m_ increased from 2.14 to 4.71 mM with increasing of the acarbose concentration from 0 to 9.99 μM (Fig. 8B). The apparent *K*_*i*_ was calculated as 54.08 μM/L.

## Discussion

The present study is the first report that LA could inhibit the feeding behaviors of herbivorous insects. Tobacco is one of the high value hosts of *H. armigera* (Fitt 1989), but our result showed tobacco leaves exposed to ≥ 10 mM LA significantly deterred the feedings of *H. armigera* caterpillars. Under both field and lab conditions, pepper leaves were not the preferred choice of *H. armgiera* caterpillars (Liu et al. 2009; Ruan and Wu 2001; Tang et al. 2006), whereas our data showed *H. armgiera* preferred control pepper leaves over leaves exposed to ≥ 10 mM LA. Sucrose and fructose are two phagostimulants of lots of lepidopteran species (Bernays and Chapman 2000; 2001; Simmonds et al. 1992; Sun et al. 2021; Vickerman and de Boer 2002), additions of sucrose or fructose on tobacco or pepper leaves could significantly induce preferences of *H. armigera* caterpillars, but the preferences could be counteracted after LA mixed to sucrose or fructose in our present study. These comprehensive behavioral results show that LA is an antifeedant compound or deterrent substance to caterpillars of the generalist herbivorous *H. armigera*.

The definition of an antifeedant could be: “a peripherally mediated behavior-modifying substance resulting in feeding deterrence” (Isman 1994; Koul 2005). In detail, antifeedants could be of pre-ingestive feeding inhibitors that act through gustatory receptors eliciting rejection of the plant material by changing the sensory code from “acceptable” to “unacceptable” (Dancewicz et al. 2020; Frazier and Chyb 1995). Our results demonstrated that the additions of LA to sucrose and tobacco leaf saps resulted in the suppressing of the sensitivities of peripheral GRNs to these substances at dose-dependent patterns, which is consistent with the tendency models of LA effects on the feeding behaviors for these substances, suggesting that the suppressions may contribute to the deterrent effects of LA on *H. armigera* caterpillars.

LA sensitive GRNs were not found in the peripheral maxillary sensillum of *H. armigra* caterpillars, indicating the indirect detection of LA in this *Helicoverpa* species. Although indirect detection of sweeteners have not yet been reported in herbivorous insects, indirect detection of deterrents by suppressing the activity of sugar GRNs have been found in *Drosophila* (Meunier et al. 2003) and the honeybee *Apis mellifera* (de Brito Sanchez et al. 2014). Our study showed the activities of GRN sensitive to phagostimulants and host leaf saps were suppressed by LA, but the mechanism of indirect detection of LA and the suppressing coding retains to be identified.

The impacts of LA on behaviors and taste could be further reflected by the prolonged larval development and the suppressed intestinal sucrase activity of *H. armgiera* caterpillars after LA ingestion. The depressed sucrase activity in midgut resulted in the accumulations of sucrose, the decreasing of glucose level and thereby the prolonged development, which is similar to the effects of prolonged duration in other insect caterpillars after digestions of toxic substances or deterrents (Glendinning 2002; Sollai et al. 2017; Yadav et al. 2010). Consequently, our results of behaviors, gustation, development and intestinal physiology provide a comprehensive understanding of the effects of LA, demonstrating that LA is an effective antifeedent to the polyphagous herbivorous insect *H. armigera*.

However, our results of dietary LA suppressing the sucrase activity is not consistent with the effects of LA on the piercing-sucking mouthpart whitefly *B. tabaci* that dietary LA exposures didn’t change the activity of α-glucosidase although LA exposure resulted in a significant drop in the mean day survival of this species (Hu et al. 2010). We postulate two possibilities may contribute to the inconsistence of α-glucosidase activity between the two insect species. Firstly, in our present experiment, we investigated the effects of LA on the intestinal sucrase activity but not the activities of all α-glucosidases because of the latter including sucrase, maltase and glucoamylase. In mammals, dietary LA showed no inhibitory effect on the activities of intestinal maltase and glucoamylase (Halschou-Jensen et al. 2015; Seri et al. 1996). But the activity of the whole α-glucosidases were investigated in the work of *B. tabaci* (Hu et al. 2010), then we suppose that the activity of total α-glucosidase in *B. tabaci* may cover the different changes of activity of different α-glucosidases, such as the sucrase and the maltase. Secondary, dietary LA may indeed have no obvious effect on the sucrase activity in *B. tabaci* because of great differences of digestive system between chewing mouthpart caterpillars and the hemipteran *B. tabaci*. In the work of *B. tabaci*, dietary LA resulted in a high mortality whereas our work showed the mortality of *H. armigera* caterpillars was not significantly changed by LA exposure.

LA is one of the components of plant biopolymers such as cereal fiber and pectin in a variety of crops. But no reports show that crops containing natural LA deter feedings of herbivorous insects, which may due to the low concentration of LA in plants (Kotake et al. 2016) and the tolerance of low concentration of deterrents in herbivorous insects (Ahn et al. 2011; Sun et al. 2019; Wink and Theile 2002). As one of plant specific primary metabolites, the metabolic pathway of LA may be of importance to plant pollen development (Kotake et al. 2016). Then its physiological functions in plants may be quite different from most reported antifeedants, such as azadirachtin, rotenone, matrine, nicotine, pyrethrins, servicing as defense compounds (Cheng et al. 2020; Isman 2020; Pavela 2016; Siegwart et al. 2015; Walia et al. 2017).

As a non-caloric sugar, the physiological and ecological roles of LA in plants is not quite clear (Kotake et al. 2016), but it is obvious that LA doesn’t service as nutrients to animals. The negative effects of LA on a variety of insects and thereby the phenotypic profiles in these insects may be different. In *Drosophila*, LA services as a peripheral sweetener, but LA exposures could be remembered as a negative experience and leaded to a so-called caloric frustration memory, resulting in a reduced response to LA, which may be involved in the post-ingestion process of this compound (Musso et al. 2017). Indeed, during the feeding processes of our work, some *H. armigera* caterpillars vomited after consumption of the LA-treated plant leaves, indicating LA evoked an emesis response of digestive disruption, which suggests the post-ingestive feedback process may also mediate the evaluation of this pentose. We are not clear whether the emesis response of *H. armgiera* caterpillars is related to the suppressed activity of intestinal sucrase in midgut, but it seems this non-caloric sugar had serious negative effects on the digestive system of caterpillars.

Together with our findings, our study demonstrated that LA could be an effective feeding antifeedant towards *H. armgiera*, depending on its inhibiting activities on caterpillars from behaviors to physiologies. We also investigated whether the antifedant effects of LA could be observed in other insect species. Indeed, our preliminary experiments suggested LA also could inhibit the feedings in another *Helicoverpa* species, the tobacco budworm *H. assulta*, and the eusocialinsect species, the termite *Reticulitermes chinensis* (Isoptera: Rhinotermitidae), but the antifeedant effect was not obvious in another extreme pest species, the fall armyworm *Spodoptera frugiperda* (J. E. Smith) (unpublished data). As a matter of fact, exploring and application of potential antifeedants would be very helpful for pest control in the current and future IPM strategy. Our work provides substantial evidence of an unusual botanical sugar which could service as a potential biological antifeedant for pest control.

## Acknowledgments

We thank Dr. Wenge Chen, Ms. Juan Qu, Dr. Huijuan Yang, and Mr. Zhongwei Sun for kindly helping in the kinetic analysis of enzyme reaction, insect rearing, supplying the tobacco seedlings, and supplying the pepper seedlings, respectively. The work was supported by the National Natural Science Foundation of China (Grant No. 31672367 & 31861133019), the Science and Technology Program of Henan Province of China (Grant No. 202102110072) and the Key Science and Technology Research Project of Henan Province of China (Grant No. 201300111500).

## Author Contributions

QT, YM, XZ, and FY designed the research. LS and ZH performed most of the experiments. YM, WH and LL performed some of the behavioral and electrophysiological experiments.

YM, LS, ZH analyzed the data. QT, YM, XY and CW wrote the manuscript. All authors read and approved the final manuscript.

## References

Ahn SJ, Badenes-Perez FR,Heckel DG (2011) A host-plant specialist, Helicoverpa assulta, is more tolerant to capsaicin from Capsicum annuum than other noctuid species. J Insect Physiol 57: 1212–1219. https://doi.org/10.1016/j.jinsphys.2011.05.015.

Akhtar Y, Yu Y, Isman MB,Plettner E (2010) Dialkoxybenzene and dialkoxyallylbenzene feeding and oviposition deterrents against the cabbage looper, Trichoplusia ni: potential insect behavior control agents. J Agric Food Chem 58: 4983–4991. https://doi.org/10.1021/jf9045123.

Bengtsson S (1990) Isolation and chemical characterization of water soluble arabinoxylans in rye grain. Carbohydr Polym 12: 267–277. https://doi.org/10.1016/0144-8617(90)90068-4.

Bernays EA,Chapman RF (2000) A neurophysiological study of sensitivity to a feeding deterrent in two sister species of Heliothis with different diet breadths. J Insect Physiol 46: 905–912. https://doi.org/10.1016/S0022-1910(99)00197-3.

Bernays EA,Chapman RF (2001) Taste cell responses in the polyphagous arctiid, Grammia geneura: towards a general pattern for caterpillars. J Insect Physiol 47: 1029–1043. https://doi.org/10.1016/s0022-1910(01)00079-8.

Cheng X, He H, Wang WX, Dong F, Zhang H, Ye J, Tan C, Wu Y, Lv X, Jiang X,Qin X (2020) Semi-synthesis and characterization of some new matrine derivatives as insecticidal agents. Pest Manag Sci 76: 2711–2719. https://doi.org/10.1002/ps.5817.

Dancewicz K, Szumny A, Wawrzenczyk C,Gabrys B (2020) Repellent and antifeedant activities of citral-derived lactones against the peach potato aphid. Int J Mol Sci 21: 8029. https://doi.org/10.3390/ijms21218029.

de Brito Sanchez MG, Lorenzo E, Su S, Liu F,Giurfa M (2014) The tarsal taste of honey bees: behavioral and electrophysiological analyses. Front Behav Neurosci 8. https://doi.org/10.3389/fnbeh.2014.00025.

DuBois GE,Prakash I (2012) Non-caloric sweeteners, sweetness modulators, and sweetener enhancers. Annu Rev Food Sci Technol 3: 353–380. https://doi.org/10.1146/annurev-food-022811-101236.

FAO (2019) New standards to curb the global spread of plant pests and diseases. Web page of the Food and Agriculture Organization of the United Nations. http://www.fao.org/news/story/en/item/1187738/icode/. Accessed 16 November, 2020.

Fitt GP (1989) The ecology of Heliothis species in relation to agroecosystems. Annu Rev Entomol 34: 17–52. https://doi.org/10.1146/annurev.en.34.010189.000313.

Frazier JL,Chyb S (1995) Use of feeding inhibitors in insect control. In: Chapman RF,de Boer G (ed) Regulatory Mechanisms in Insect Feeding, Chapman & Hall, New York, NY, USA, pp. 364–381.

Garleb KA, Bourquin LD,Fahey GC (1989) Neutral monosaccharide composition of various fibrous substrates: a comparison of hydrolytic procedures and use of anion-exchange high-performance liquid-chromatography with pulsed amperometric detection of monosaccharides. J Agric Food Chem 37: 1287–1293. https://doi.org/10.1021/jf00089a018.

Glendinning JI (2002) How do herbivorous insects cope with noxious secondary plant compounds in their diet? Entomol Exp Appl 104: 15–25. https://doi.org/10.1046/j.1570-7458.2002.00986.x.

Halschou-Jensen K, Bach Knudsen KE, Nielsen S, Bukhave K,Andersen JR (2015) A mixed diet supplemented with L-arabinose does not alter glycaemic or insulinaemic responses in healthy human subjects. Br J Nutr 113: 82–88. https://doi.org/10.1017/S0007114514003407.

Halschoujensen K, Knudsen KE, Nielsen S, Bukhave K,Andersen JR (2015) A mixed diet supplemented with L-arabinose does not alter glycaemic or insulinaemic responses in healthy human subjects. Br J Nutr 113: 82–88. https://doi.org/10.1017/S0007114514003407.

Hasheminia SM, Sendi JJ, Jahromi KT,Moharramipour S (2013) Effect of milk thistle, Silybium marianum, extract on toxicity, development, nutrition, and enzyme activities of the small white butterfly, Pieris rapae. J Insect Sci 13: 146. https://doi.org/10.1673/031.013.14601.

Hu JS, Gelman DB, Salvucci ME, Chen YP,Blackburn MB (2010) Insecticidal activity of some reducing sugars against the sweet potato whitefly, Bemisia tabaci, Biotype B. J Insect Sci 10: 203. https://doi.org/10.1673/031.010.20301.

Isman MB (1994) Botanical insecticides and antifeedants: new sources and perspectives. Pestic Res J 6: 11–19.

Isman MB (2020) Commercial development of plant essential oils and their constituents as active ingredients in bioinsecticides. Phytochem Rev 19: 235–241. https://doi.org/10.1007/s11101-019-09653-9.

Jiang J, Ding S, Zhang Y,Wang Y (2010) Effects of artificial diet on the growth & development and fecundity of H. armigera. J Henan Agr Univ 44: 78–82. https://doi.org/10.16445/j.cnki.1000-2340.2010.01.007.

Jiang X, Xu H, Zheng N, Yin X,Zhang L (2020) A Chemosensory Protein Detects Antifeedant in Locust (Locusta migratoria). Insects 12: 1. https://doi.org/10.3390/insects12010001.

Junhirun P, Pluempanupat W, Yooboon T, Ruttanaphan T, Koul O,Bullangpoti V (2018) The study of isolated alkane compounds and crude extracts from Sphagneticola trilobata (Asterales: Asteraceae) as a candidate botanical insecticide for lepidopteran larvae. J Econ Entomol 111: 2699–2705. https://doi.org/10.1093/jee/toy246.

Karkanis AC,Athanassiou CG (2021) Natural insecticides from native plants of the Mediterranean basin and their activity for the control of major insect pests in vegetable crops: shifting from the past to the future. J Pest Sci 94: 187–202. https://doi.org/10.1007/s10340-020-01275-x.

Kotake T, Yamanashi Y, Imaizumi C,Tsumuraya Y (2016) Metabolism of L-arabinose in plants. J Plant Res 129: 1–12. https://doi.org/10.1007/s10265-016-0834-z.

Koul O (2005) Insect Antifeedants. CRC Press, Boca Raton, USA.

Koul O, Daniewski WM, Multani JS, Gumulka M,Singh G (2003) Antifeedant effects of the limonoids from Entandrophragma candolei (Meliaceae) on the gram pod borer, Helicoverpa armigera (Lepidoptera: Noctuidae). J Agric Food Chem 51: 7271–7275. https://doi.org/10.1021/jf0304223.

Liu Z, Gong P, Heckel DG, Wei W, Sun J,Li D (2009) Effects of larval host plants on over-wintering physiological dynamics and survival of the cotton bollworm, Helicoverpa armigera (Hubner) (Lepidoptera: Noctuidae). J Insect Physiol 55: 1–9. https://doi.org/10.1016/j.jinsphys.2008.07.017.

Ma Y, Li JJ, Tang QB, Zhang XN, Zhao XC, Yan FM,van Loon JJA (2016) Trans-generational desensitization and within-generational resensitization of a sucrose-best neuron in the polyphagous herbivore Helicoverpa armigera (Lepidoptera: Noctuidae). Sci Rep 6: 39358. https://doi.org/10.1038/srep39358.

Mao G, Tian Y, Sun Z, Ou J,Xu H (2019) Bruceine D isolated from Brucea Javanica (L.) Merr. as a systemic feeding deterrent for three major lepidopteran pests. J Agric Food Chem 67: 4232–4239. https://doi.org/10.1021/acs.jafc.8b06511.

Meunier N, Marion-Poll F, Rospars JP,Tanimura T (2003) Peripheral coding of bitter taste in Drosophila. J Neurobiol 56: 139–152. https://doi.org/10.1002/neu.10235.

Mitri C, Soustelle L, Framery B, Bockaert J, Parmentier ML,Grau Y (2009) Plant insecticide L-canavanine repels Drosophila via the insect orphan GPCR DmX. PLoS Biol 7: e1000147. https://doi.org/10.1371/journal.pbio.1000147.

Musso P, Lampin-Saint-Amaux A, Tchenio P,Preat T (2017) Ingestion of artificial sweeteners leads to caloric frustration memory in Drosophila. Nat Commun 8: 1803. https://doi.org/10.1038/s41467-017-01989-0.

Pavela R (2016) History, presence and perspective of using plant extracts as commercial botanical insecticides and farm products for protection against insects - a review. Plant Prot Sci 52: 229–241. https://doi.org/10.17221/31/2016-PPS.

Ponsankar A, Sahayaraj K, Senthil-Nathan S, Vasantha-Srinivasan P, Karthi S, Thanigaivel A, Petchidurai G, Madasamy M,Hunter WB (2020) Toxicity and developmental effect of cucurbitacin E from Citrullus colocynthis L. (Cucurbitales: Cucurbitaceae) against Spodoptera litura Fab. and a non-target earthworm Eisenia fetida Savigny. Environ Sci Pollut Res Int 27: 23390–23401. https://doi.org/10.1007/s11356-019-04438-1.

Roessingh P, Hora KH, van Loon JJA,Menken SBJ (1999) Evolution of gustatory sensitivity in Yponomeuta caterpillars: sensitivity to the stereo-isomers dulcitol and sorbitol is localised in a single sensory cell. J Comp Physiol A 184: 119–126. https://doi.org/10.1007/s003590050311.

Ruan YM,Wu KJ (2001) Performances of the cotton bollworm, Helicoverpa armigera on different food plants. Acta Entomol Sin 44: 205–212. https://doi.org/10.16380/j.kcxb.2001.02.013.

Saulnier L, Vigouroux J,Thibault JF (1995) Isolation and partial characterization of feruloylated oligosaccharides from maize bran. Carbohydr Res 272: 241–253. https://doi.org/10.1016/0008-6215(95)00053-V.

Seri K, Sanai K, Matsuo N, Kawakubo K, Xue C,Inoue S (1996) L-arabinose selectively inhibits intestinal sucrase in an uncompetitive manner and suppresses glycemic response after sucrose ingestion in animals. Metabolism 45: 1368–1374. https://doi.org/10.1016/S0026-0495(96)90117-1.

Shibanuma K,Houda K (2011) Determination of the transient period of the EIS complex and investigation of the suppression of blood glucose levels by L-arabinose in healthy adults. Eur J Nutr 50: 447–453. https://doi.org/10.1007/s00394-010-0154-3.

Siegwart M, Graillot B, Lopez CB, Besse S, Bardin M, Nicot PC,Lopez-Ferber M (2015) Resistance to bio-insecticides or how to enhance their sustainability: a review. Front Plant Sci 6. https://doi.org/10.3389/Fpls.2015.00381.

Simmonds MS, Simpson SJ,Blaney WM (1992) Dietary selection Behaviour in Spodoptera Littoralis: the effects of conditioning diet and conditioning period on neural responsiveness and selection behaviour. J Exp Biol 162: 73–90.

Sollai G, Biolchini M, Solari P,Crnjar R (2017) Chemosensory basis of larval performance of Papilio hospiton on different host plants. J Insect Physiol 99: 47–57. https://doi.org/10.1016/j.jinsphys.2017.02.007.

Sollai G, Tomassini Barbarossa I, Masala C, Solari P,Crnjar R (2014) Gustatory sensitivity and food acceptance in two phylogenetically closely related papilionid species: Papilio hospiton and Papilio machaon. PLoS One 9: e100675. https://doi.org/10.1371/journal.pone.0100675.

Sollai G, Tomassini Barbarossa I, Solari P,Crnjar R (2015) Taste discriminating capability to different bitter compounds by the larval styloconic sensilla in the insect herbivore Papilio hospiton (Gene). J Insect Physiol 74: 45–55. https://doi.org/10.1016/j.jinsphys.2015.02.004.

Sun L, Hou W, Zhang J, Dang Y, Yang Q, Ma Y,Tang Q (2021) Plant metabolites drive different responses in caterpillars of two closely related Helicoverpa species. Front Physiol. https://doi.org/10.3389/fphys.2021.662978.

Sun Z, Shi Q, Li Q, Wang R, Xu C, Wang H, Ran C, Song Y,Zeng R (2019) Identification of a cytochrome P450 CYP6AB60 gene associated with tolerance to multi-plant allelochemicals from a polyphagous caterpillar tobacco cutworm (Spodoptera litura). Pestic Biochem Physiol 154: 60–66. https://doi.org/10.1016/j.pestbp.2018.12.006.

Tang Q, Huang L, Wang C, Zhan H,van Loon JJA (2014) Inheritance of electrophysiological responses to leaf saps of host-and nonhost plants in two Helicoverpa species and their hybrids. Arch Insect Biochem Physiol 86: 19–32. https://doi.org/10.1002/arch.21154.

Tang Q, Jiang J, Yan Y, van Loon JJA,Wang C (2006) Genetic analysis of larval host-plant preference in two sibling species of Helicoverpa. Entomol Exp Appl 118: 221–228. https://doi.org/10.1111/j.1570-7458.2006.00387.x.

van Loon JJA (1990) Chemoreception of phenolic acids and flavonoids in larvae of two species of Pieris. J Comp Physiol A 166: 889–899. https://doi.org/10.1007/bf00187336.

van Opstal AM, Kaal I, van den Berg-Huysmans AA, Hoeksma M, Blonk C, Pijl H, Rombouts S,van der Grond J (2019) Dietary sugars and non-caloric sweeteners elicit different homeostatic and hedonic responses in the brain. Nutrition 60: 80–86. https://doi.org/10.1016/j.nut.2018.09.004.

Vickerman DB,de Boer G (2002) Maintenance of narrow diet breadth in the monarch butterfly caterpillar: response to various plant species and chemicals. Entomol Exp Appl 104: 255–269. https://doi.org/DOI10.1046/j.1570-7458.2002.01012.x.

Wada-Katsumata A, Silverman J,Schal C (2013) Changes in taste neurons support the emergence of an adaptive behavior in cockroaches. Science 340: 972–975. https://doi.org/10.1126/science.1234854.

Walia S, Saha S, Tripathi V,Sharma KK (2017) Phytochemical biopesticides: some recent developments. Phytochem Rev 16: 989–1007. https://doi.org/10.1007/s11101-017-9512-6.

Wang Y, Ma Y, Zhou D, Gao S, Zhao X, Tang Q, Wang C,van Loon JJA (2017) Higher plasticity in feeding preference of a generalist than a specialist: experiments with two closely related Helicoverpa species. Sci Rep 7: 17876. https://doi.org/10.1038/s41598-017-18244-7.

Wink M,Theile V (2002) Alkaloid tolerance in Manduca sexta and phylogenetically related sphingids (Lepidoptera : Sphingidae). Chemoecology 12: 29–46. https://doi.org/10.1007/s00049-002-8324-2.

Wroblewska-Kurdyk A, Dancewicz K, Gliszczynska A,Gabrys B (2020) New insight into the behaviour modifying activity of two natural sesquiterpenoids farnesol and nerolidol towards Myzus persicae (Sulzer) (Homoptera: Aphididae). Bull Entomol Res 110: 249–258. https://doi.org/10.1017/S0007485319000609.

Yadav J, Tan CW,Hwang SY (2010) Spatial variation in foliar chemicals within radish (Raphanus sativus) plants and their effects on performance of Spodoptera litura. Environ. Entomol. 39: 1990–1996. https://doi.org/10.1603/EN10118.

Yamamoto H (2013) Physiological functions and food application of L-arabinose. Oleoscience 13: 429–434. https://doi.org/10.5650/oleoscience.13.429.

Zalucki MP, Daglish G, Firempong S,Twine PH (1986) The biology and ecology of Heliothis armigera (Hübner) and H. punctigera Wallengren (Lepidoptera:Noctuidae) in Australia - What do we know? Aust J Zool 34: 779–814. https://doi.org/10.1071/ZO9860779.

Zhou D, van Loon JJ,Wang CZ (2010) Experience-based behavioral and chemosensory changes in the generalist insect herbivore Helicoverpa armigera exposed to two deterrent plant chemicals. J Comp Physiol A Neuroethol Sens Neural Behav Physiol 196: 791–799. https://doi.org/10.1007/s00359-010-0558-9.

